# Lack of detection of SARS-CoV-2 in Wildlife from Kerala, India in 2020-21

**DOI:** 10.1101/2023.07.03.547244

**Authors:** Arun Zachariah, Sajesh P Krishnankutty, Jishnu Manazhi, Vishnu Omanakuttan, Sam Santhosh, Adam Blanchard, Rachael Tarlinton

## Abstract

Spill over of SARs-CoV-2 into a variety of wild and domestic animals has been an ongoing feature of the human pandemic. The establishment of a new reservoir in white tailed deer in North America and increasing divergence of the viruses circulating in them from those circulating in the human population has highlighted the ongoing risk this poses for global health. Some parts of the world have seen more intensive monitoring of wildlife species for SARS-CoV-2 and related coronaviruses but there are still very large gaps in geographical and species-specific information. This paper reports negative results for SARS-CoV-2 PCR based testing using a pan coronavirus end point RDRP PCR and a Sarbecovirus specific E gene qPCR on lung and or gut tissue from wildlife from the Indian State of Kerala. These animals included: 121 *Rhinolophus rouxii* (Rufous Horsehoe Bat), *6 Rhinolophus bedommei* (Lesser Woolly Horseshoe Bat), *15 Rossettus leschenaultii* (Fulvous Fruit Bat), *47 Macaca radiata* (Bonnet macaques), *35 Paradoxurus hermaphroditus (*Common Palm Civet), *5 Viverricula indica* (Small Indian Civet), *4 Herpestes edwardsii* (Common Mongoose), *10 Panthera tigris* (Bengal Tiger), *8 Panthera pardus fusca* (Indian Leopard), 4 *Prionailurus bengalensis* (Leopard cats), 2 *Felis chaus* (Jungle cats), 2 *Cuon alpinus* (Wild dogs) and 1 *Melursus ursinus* (sloth bear).

## 3. Introduction

There have been numerous reports of SARS-CoV-2 spill over from the human pandemic into multiple species. Prominent events with large numbers of animals in multiple sites and spill over back into the human population include domestic cats (*Felis Cattus)* (Piewbang, Poonsin et al. 2022, Tyson, Jones et al. 2023), farmed American mink (*Neogale vison)* (Wasniewski, Boué et al. 2023) (Oude Munnink, Sikkema et al. 2021) and Syrian hamsters (*Mesocricerus auratus*) (Kok, Wong et al. 2022, Yen, Sit et al. 2022). SARS-CoV-2 has also established ongoing transmission in wild white-tailed deer (*Odocoileus virginianus*) in the USA, with infection back into the human population confirmed. Worryingly the variants found in the deer population have begun to significantly diverge from those in the human population creating an unpredictable reservoir of novel variants (Pickering, Lung et al. 2022, Caserta, Martins et al. 2023, McBride, Garushyants et al. 2023). It would also appear from laboratory studies that the range of species able to be infected by SARS-CoV-2 is very dependent on the strain of virus and it is likely that as it continues to evolve in people that the species range of susceptibility will not be stable (Halfmann, Iida et al. 2022, Thakur, Gallo et al. 2022).

There have been a very large number of reports of other species either able to be infected experimentally or with infection detected in sporadic case reports. These are reviewed in (Nielsen, Alvarez et al. 2023) but include a large number of cricetid rodents, felids, mustelids, other small carnivores and primates. Many of these reports have been from animals held in zoological collections where they are in close contact with humans, and it is not clear whether these species in their natural environment are at risk or not. Indeed there is a marked contrast in disease transmission between farmed mink at high population density, with almost 100% of animals infected in a very short period of time in some outbreaks (Boklund, Hammer et al. 2021) and the sporadic reports, despite intense monitoring, in wild animals, which are largely solitary (Aguiló-Gisbert, Padilla-Blanco et al. 2021, Shriner, Ellis et al. 2021, Sikkema, Begeman et al. 2022, Villanueva-Saz, Giner et al. 2022).

These behavioural considerations may be as important as biological barriers in which species the virus establishes in. In addition, we also still have very large gaps in knowledge of the distribution of sarbecoviruses in bats from the Rhinolophoidea; horseshoe bats and roundleaf bats, their natural hosts. There has been intensive sampling of bats in SE Asia, driven by the original SARS-CoV outbreak in 2006 (Wong, Li et al. 2019). This effort has established that Sarbecoviruses (SARS like betacoronaviruses) are largely only found in Rhinolophoidid bats. There are however about 180 species of these bats spread across Eurasia and Africa with coronaviruses detected in about 30 of them (Muylaert, Kingston et al. 2022). Central and South Asia alongside Sub-Saharan Africa are notable absences in *Sarbecovirus* detection studies (Cohen, Fagre et al. 2023) with that gap only just beginning to be filled (Geldenhuys, Mortlock et al. 2021, Kamau, Ergunay et al. 2022, Kettenburg, Kistler et al. 2022, Ntumvi, Ndze et al. 2022, Meta Djomsi, Lacroix et al. 2023).

There has been remarkably little study of SARS-CoV-2 in animals in India despite the countries devastating human pandemic (Wani, Menon et al. 2023). One study in Gujarat (a north western state) of 413 domestic animals of a variety of species reported 23.79% of animals qPCR positive on nasal or rectal swabs, the positive animals being dogs, cattle and buffalo with sequence confirmation of one canine isolate (Kumar, Antiya et al. 2022). Sequencing effort was targeted in areas with large number of human cases potentially explaining the very high qPCR positivity in this study. A serological study of 320 captive Bengal tigers, Asiatic lions and leopards from 8 Indian states demonstrated that 48 (15%) of these animals had seroconverted to SARS-CoV-2 by October 2021. A small number of Indian Elephants (24) and 40 spotted and swamp deer were all seronegative (Borkakoti, Karikalan et al. 2023). There have also been reports of PCR positive Asiatic lions in zoos (Karikalan, Chander et al. 2022) with 2/18 animals in Uttar Pradesh (Northern India) and 1/20 in Rajasthan (North West India) qPCR positive on nasal or rectal swabs, with sequence confirmation of the isolates, other felids housed at these institutions did not test positive. Four out of 24 Asiatic lions in Chennai (Tamil Nadu state, South East India) were also found to be qPCR positive and sequence confirmed in a zoo (Mishra, Kumar et al. 2021), two of these animals died. The only report in a wild animal in India is a solitary juvenile Asiatic leopard found dead in Uttar Pradesh with qPCR positivity and sequence confirmation in (Mahajan, Karikalan et al. 2022), this was the only animal out of more than 500 qPCR screened samples positive. In all these cases the felid infections were consistent with the circulating human variants at the time.

India’s size and number of climate zones mean that biodiversity is very high with pressures from the world’s largest human population and known problems with illegal wildlife trade and human/wildlife conflict contributing to multiple zoonotic disease outbreaks (Walsh and Hossain 2020, Goodale, Mammides et al. 2022, Rana and Kumar 2023). The western ghats rainforest along the west coast of India is a biodiversity hotspot with 133 mammal species recorded. It is also an area of intense human wildlife interaction and conflict, with large species such as tigers and elephants causing considerable destruction in human settlements. Consequent to this zoonotic disease outbreaks are frequent with the Kyasanur forest and its eponymous virus part of this ecosystem. Surveillance systems and monitoring in this region are however seriously under-resourced with little systematic surveillance of either animals or their viruses (Walsh and Hossain 2020).

This study sought to partially bridge these gaps with targeted trapping and testing of Rhinolophus bats and opportunistic testing of carnivore and primate species either found dead (road kill) or culled as part of nuisance animal control activities in the state of Kerala in South West India.

## 4. Methods

### 4.1 Sample collection

A total of 260 animals from 13 species (Table 1) were targeted for coronavirus monitoring. For the two horseshoe bat species, palm civets and common mongoose, the targeted numbers were calculated in Epitools (Epitools 2020), 2 stage sampling for demonstration of disease freedom (cluster size unknown) based on assumption of 5% prevalence of Coronavirus and 50% of populations affected. Prior assumptions were based on previous studies of rodent coronaviruses in wild populations (Tsoleridis and Ball 2020). This gave an estimate of 7 clusters with 17 individuals in each cluster to be samples (119 animals per species). The species targeted were the two most common horseshoe bats in this environment (others are rare) and the most common small carnivore predators of bats in these sites.

**Table 1:**
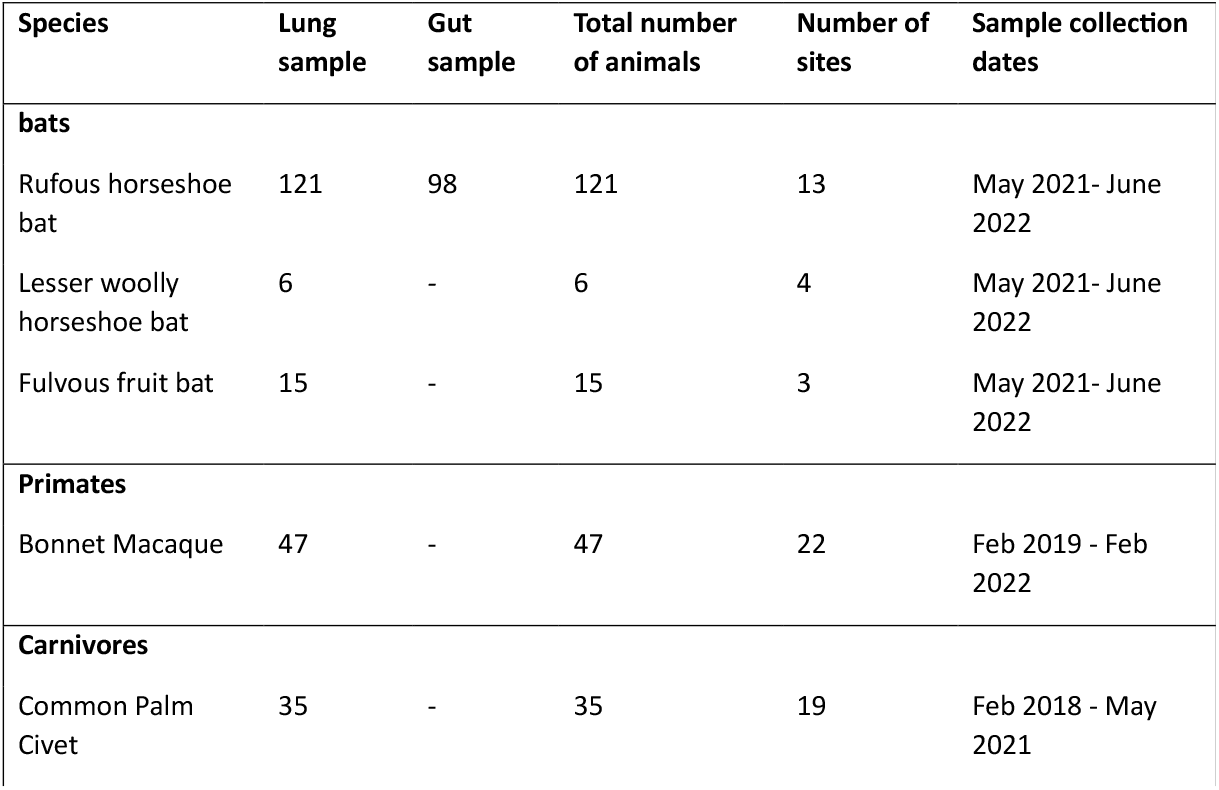
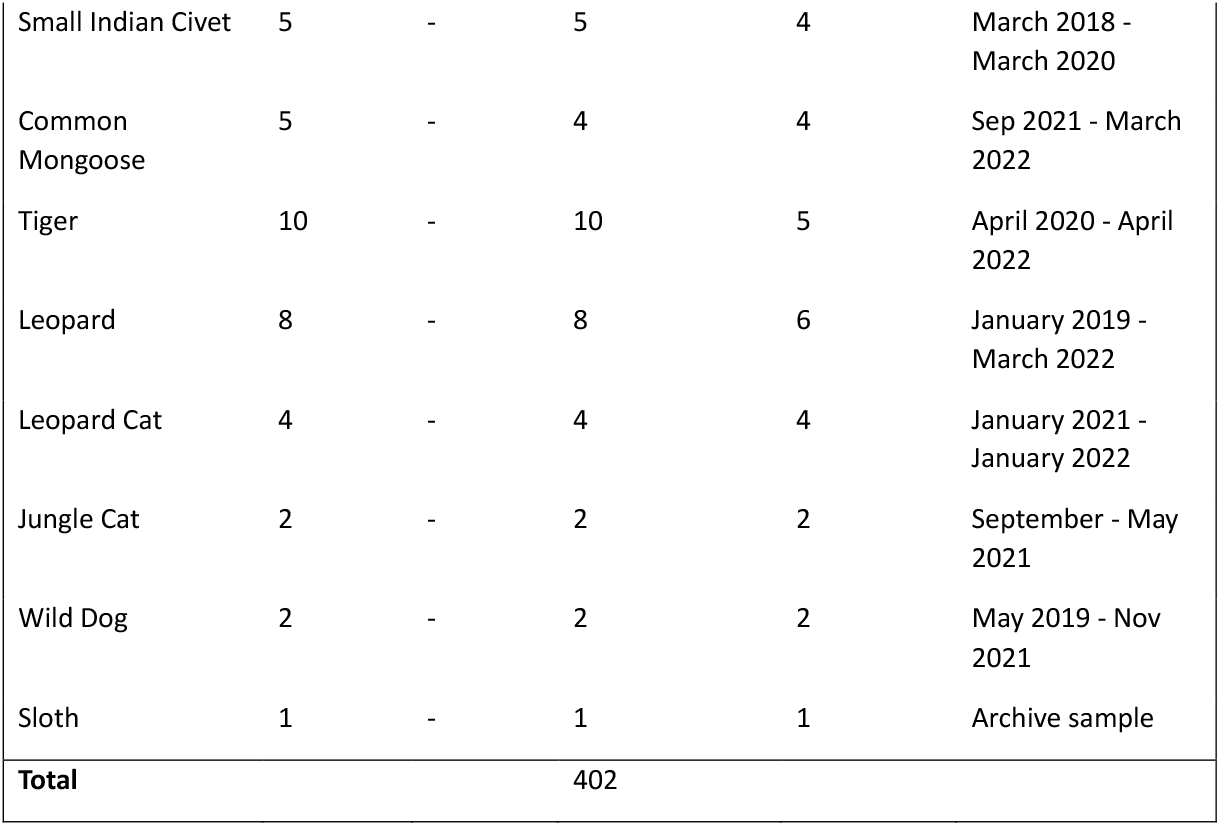
Species and Sample Type Screened for SARS-CoV-2

| Species | Lung sample | Gut sample | Total number of animals | Number of sites | Sample collection dates |
| --- | --- | --- | --- | --- | --- |
| **bats** |  |  |  |  |  |
| Rufous horseshoe bat | 121 | 98 | 121 | 13 | May 2021- June 2022 |
| Lesser woolly horseshoe bat | 6 | - | 6 | 4 | May 2021- June 2022 |
| Fulvous fruit bat | 15 | - | 15 | 3 | May 2021- June 2022 |
| **Primates** |  |  |  |  |  |
| Bonnet Macaque | 47 | - | 47 | 22 | Feb 2019 - Feb 2022 |
| **Carnivores** |  |  |  |  |  |
| Common Palm Civet | 35 | - | 35 | 19 | Feb 2018 - May 2021 |
| Small Indian Civet | 5 | - | 5 | 4 | March 2018 -<br>March 2020 |
| Common<br>Mongoose | 5 | - | 4 | 4 | Sep 2021 - March<br>2022 |
| Tiger | 10 | - | 10 | 5 | April 2020 - April<br>2022 |
| Leopard | 8 | - | 8 | 6 | January 2019 -<br>March 2022 |
| Leopard Cat | 4 | - | 4 | 4 | January 2021 -<br>January 2022 |
| Jungle Cat | 2 | - | 2 | 2 | September - May<br>2021 |
| Wild Dog | 2 | - | 2 | 2 | May 2019 - Nov<br>2021 |
| Sloth | 1 | - | 1 | 1 | Archive sample |
| **Total** |  |  | 402 |  |  |

Bats were trapped using mist or harp nets, Subject to inhalational anaesthesia with isoflurane with throat and cloacal swabs collected. Later bats were euthanized by extending the anaesthesia and tissue samples were collected by necropsy. Samples were stored in RNAlater for nucleic acid extraction. Small carnivores (common palm civet, small Indian civet, common mongoose, leopard cat, jungle cat), bonnet macaques and larger carnivores (Bengal tiger, leopard, sloth bear and wild dog), samples were collected as part of routine necropsy procedures from dead animals in the study area. All the carcasses were fresh (within 12 hours of death) and samples were preserved in RNA later and stored in -80 degree Celsius. All procedures were conducted under the supervision of an experienced wildlife veterinarian.

Ethical approval was granted by the University of Nottingham School of Veterinary Medicine and Science Committee for Animal Research and Ethics (CARE). Permission for Field work in Forest Areas for Scientific Research and sample collection was as per the permit number KFDHQ-1979/ 2021-CWW / WL 10 issued by the Chief Wildlife Warden, Kerala state, India.

### 4.2 RNA extraction, reverse transcriptase (RT) and RNA-dependent RNA polymerase (RDRP) gene coronaviruses generic conventional PCR

All sample processing and PCR was performed in India at the Kerala State forest department and SciGenom labs, Kerala.

RNA extraction from lung tissue, faecal samples, rectal and oronasal swabs, and cell culture supernatant as positive control, was carried out using the Invitrogen Viral RNA extraction kit as per manufacturer’s instructions. The positive control sample used throughout this study was cDNA from the OC43 Coronavirus ATCC strain VR1558). RT was performed with the Applied Biosystems cDNA reverse transcription kit as per manufacturer’s instructions. All cDNA products were stored at -20 °C for conventional PCR. An endpoint SARS-CoV-2 Specific PCR assay (Poon, Chu et al. 2005, Tsoleridis and Ball 2020) was used to amplify the RDRP gene with the Takara R050 A PrimeSTAR GXL taq according to manufacturer’s instructions.

RNA and cDNA quality control was assessed via partial amplification of 108 bp of the beta actin gene using a published conventional PCR protocol (Fischer, Freuling et al. 2014). Primers were F: CAGCACAATGAAGATCAAGATCATC and R: CGGACTCATCGTACTCCTGCTT

## 5. Results

No animal sample tested positive for SARS-CoV-2. Locations of samples are shown in Figure 1.

**Figure 1.**
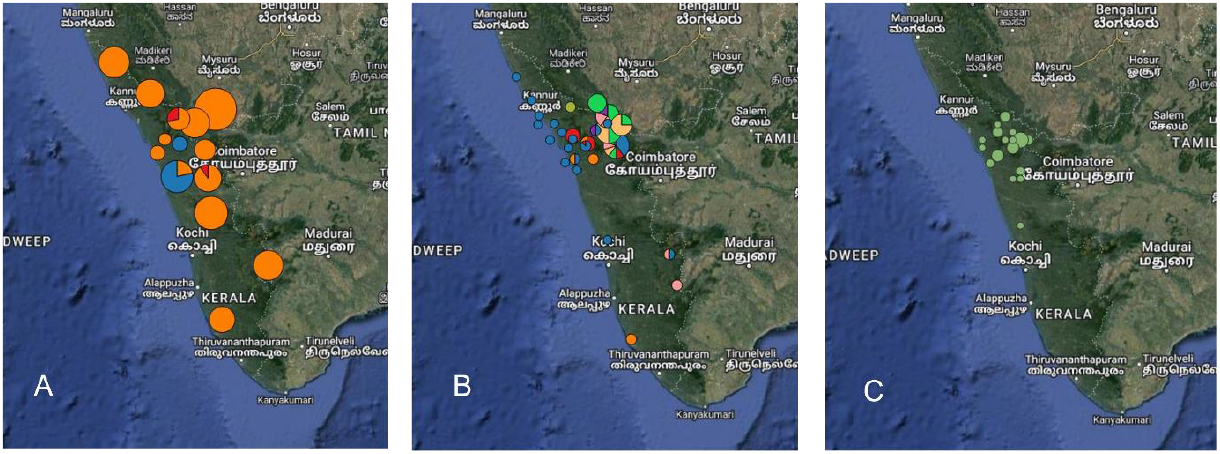
a) locations of bat samples, orange= rufous horseshoe bad red = lessor woolly horseshoe bat, blue = fulvous fruit bat, pie chart sizes are proportional to the number of animals at each location b) locations of carnivore samples, blue=common palm civet, red=small Indian civet, orange = common mongoose, bright green=tiger, yellow=leopard, pink = leopard cat, purple= jungle cat, light green = wild dog, grey= sloth bear pie chart sizes are proportional to the number of animals at each location, c) locations of bonnet macaque samples, circles are proportional to the number of animals at each location (Maps drawn in QGIS v 3.3.1)

## 6. Discussion

This study found no evidence of widespread circulation of SARs-CoV-2 or related coronaviruses in Indian wildlife. Some of the species tested here, such as bonnet macaques, palm civets and mongoose are very commonly found in and around human habitation and represent significant pest or nuisance species in terms of aggressive interactions with humans and potential zoonoses or cross species transmission to and from domestic animals (Sato, Kabeya et al. 2013, Balasubramaniam, Marty et al. 2020, Kadam, Karikalan et al. 2022). These species are high risk for SARS-Cov-2 spill over and it is at least reassuring that these animals tested negative. Though with the large caveats that sampling was PCR based, a small number of animals and could easily have missed infections. Follow up work with serological testing for SARS-CoV-2 antibody (indicating previous infection) would be an extremely useful follow up to this project, with of course the caveat that widely available serological assays have not been validated for these species, making results difficult to interpret (Borkakoti, Karikalan et al. 2023).

Studies of felids in zoo (captive) populations in India have demonstrated a high rate of seroconversion (Borkakoti, Karikalan et al. 2023) and PCR positive animals have been detected in zoos (Mishra, Kumar et al. 2021, Karikalan, Chander et al. 2022) and in one wild leopard (Mahajan, Karikalan et al. 2022). Our results here, while a small number of opportunistic samples add to evidence that SARS-CoV-2 is not a widespread issue in Wild Indian Felids (Mahajan, Karikalan et al. 2022).

A completely negative finding in the two horseshoe bat species was unexpected, particularly as these species are the natural hosts of SARs-like viruses and the PCR assays used in this study should have detected known horseshoe bat sarbecoviruses. Our similar study of UK horseshoe bats did however demonstrate that presence or absence of sarbecoviruses can be very species specific with lesser horseshoe bats having a 44 % positivity rate on faecal or rectal swab samples but no detection at all in greater horseshoe bats (Apaa, Withers et al. 2023). Studies in SE Asia present with very different results with high positivity rates and sarbecoviruses detected in multiple species (Wu, Han et al. 2022). Of note the species in which SARS-CoV-2 like sarbecoviruses and recombinant viruses are commonly found *R. sinicus, R. ferrumequinum, R. pusillus, and R. affinis* are either rare (*R. pusillus*) or not found in Kerala. These species are all cave roosting bats that form large colonies which may be a key factor in facilitating sarbecovirus diversity and cross species transmission.

Our sampling numbers and targets should have been able to detect sarbecoviruses in rufous horseshoe bats where target numbers were achieved. Target numbers were not achieved in other species, primarily due to extreme adverse weather conditions (flooding) in Kerala during the sampling period. Most known roost sites for the lesser woolly horseshoe bat (which frequently roosts in sites such as drain coverts) were found abandoned. Trapping success rates for small carnivores were also less than optimal. Nonetheless we present our negative results in the interest of providing the only data to date on Indian horseshoe bat populations. This adds to data indicating that sarbecovirus spill-over out of the horseshoe bat population may be a distinctly regional (SE Asian) phenomena (Muylaert, Kingston et al. 2022, Wu, Han et al. 2022).

## Supporting information

Sample details

## 7. Author statements

### 7.3 Ethical approval

Ethical approval was granted by the University of Nottingham School of Veterinary Medicine and Science Committee for Animal Research and Ethics (CARE), and the University of Sussex Animal Welfare and Ethical Review Board.

## 7.4 Acknowledgements

